# Building a high-quality reference genome assembly for the the eastern Mediterranean Sea invasive sprinter *Lagocephalus sceleratus* (Tetraodontiformes, Tetraodontidae)

**DOI:** 10.1101/2020.02.17.952580

**Authors:** Theodoros Danis, Alexandros Tsakogiannis, Jon B. Kristoffersen, Daniel Golani, Dimitris Tsaparis, Panagiotis Kasapidis, Georgios Kotoulas, Antonios Magoulas, Costas S. Tsigenopoulos, Tereza Manousaki

## Abstract

The Tetraodontidae family encompasses several species which attract scientific interest in terms of their ecology and evolution. However, the genomic resources and especially reference assemblies are sparse for the members of the family. In this study, we focus on the silver-cheeked toadfish (*Lagocephalus sceleratus*) a well-known ‘invasive sprinter’ that has invaded and spread throughout the Eastern and part of the Western Mediterranean Sea from the Red Sea through the Suez Canal within a decade. We sequenced the genome of *L. sceleratus* using a single MinION flow cell for the main assembly, and Illumina reads for polishing the assembly. The resulted assembly consisted of 241 contigs (N50 = 11,3 Mb) with a total size of 360 Mb and yielded 98% BUSCO completeness. The high-quality genome assembly built here is expected to set the ground for future studies on this focal species’ invasive biology.

## BACKGROUND

The Suez Canal’s opening in 1869 initiated a process of invasion from the Red Sea into the Mediterranean, the so-called Lessepsian migration (Golani 2010). This influx of marine organisms has greatly impacted the local communities in ecological, evolutionary (Sax et al. 2007), and economical terms (Arim et al. 2006). Lessepsian fish comprise nowadays a significant percentage of all recorded invasive species in the Mediterranean Sea (Zenetos et al. 2012) and are under suspicion for several indigenous species displacements (Golani 2010). Lessepsian migration, having clear human driven, direct and indirect origins, is a phenomenon suitable for studying fast evolutionary change (Palumbi 2001). Genome-wide data exploration is a major process to investigate potential adaptive changes that affect invasion success.

The Silver-cheeked toadfish, *Lagocephalus sceleratus* (Gmelin 1789), is a member of the Tetraodontidae family (called puffers), widely distributed throughout the Indian and Pacific Oceans (Akyol et al. 2005). The first record of *L. sceleratus* invasion in the Mediterranean Sea, was reported in the Gökova Bay, in the south-eastern Aegean Sea coast of Turkey (Filiz and Er, 2004), and two years later in the Cretan Sea (Kasapidis et al. 2007).

Fatal toxicity, capability of fast spreading throughout the entire Levant, Aegean and Ionian Seas (Akyol & Ünal, 2017; Kalogirou, 2013), reduction of important commercial cephalopod species stocks and damaging of fishing gears (Bakiu and Durmisshaj 2019) render *L. sceleratus* one of the most significant alien fish (Streftaris and Zenetos 2006). However, the lack of a high-quality reference genome assembly hampers a genome-wide exploration of potential adaptive changes that affect its invasion success.

Following the recent advances of molecular biology and bioinformatic methodologies, as well as of sequencing technologies, the aim of this paper is to provide the first high-quality genome assembly of *L.sceleratus*, which was constructed by the combination of short but accurate Illumina reads with long but error-prone Oxford Nanopore Technology (ONT) reads. This valuable and robust genome source of *L. sceleratus*, enables future studies on ecological, evolutionary and other aspects of the species biology.

## METHODS

### 1. Sample collection, libraries construction & sequencing

Animal care and handling were carried out following well established guidelines [Guidelines for the treatment of animals in behavioral research and teaching. Anim. Behav. 53, 229–234 (1997)].

One female fish (58 cm in length) was caught alive in Hersonissos, Agios Georgios (35°20’07.50”N 25°23’11.30”E) at the pre-spawning/spawning stage (stereoscopic investigation of the oocytes) and was anesthetized using clove oil. In total, 10 mL of blood was collected using a sterilized syringe and stored in tubes that contained ∼1/10 of volume heparin for subsequent DNA extraction.

DNA extraction for the purpose of ONT sequencing was conducted on the day of sampling, from 2 μl of the freshly taken blood, using Qiagen Genomic tip (20G) and following the manufacturer’s instructions. The final elution was made with 50 μl AE buffer providing 90,4 ng/μl (Qubit measurement) of high molecular weight DNA with extra purity (Purity rates measured with Nanodrop: 260/280 = 1,87 & 260/230 = 2,12). Then, we constructed four ligation libraries (SQK-LSK109) following the manufacturer’s instructions (ref). Approximately 1.2 μg of unsheared DNA was used for each library. Two of the prepared libraries were divided into two aliquots. Each library was run for approximately 24 hours on the HCMR MinION sequencer, after which the ONT nuclease flush protocol was performed and a fresh library or library aliquot was loaded onto the same R9.4.1 flow cell. The total run time was ∼130 hours. Basecalling was done with Guppy v3.2.4 in High Accuracy Mode.

For the purpose of Illumina sequencing, we proceeded with two-days old refrigerated blood sample using the same procedure and protocol. We used 4 μl of blood eluted in 100 μl AE buffer which resulted in 79,2 ng/μl (Qubit measurement) of extra pure DNA (260/280 = 1,85 & 260/230 = 2,21).

DNA integrity was assessed by electrophoresis in 0.4 % w/v megabase agarose gel. Template DNA for Illumina sequencing was sheared by ultrasonication in a Covaris instrument. A PCR-free library was prepared with the Kapa Hyper Prep DNA kit with TruSeq Unique Dual Indexing. Paired end 2×150 bp sequencing was performed at the Norwegian Sequencing Centre (NSC) on an Illumina Hiseq4000 platform.

### 2. Data pre-processing and Genome size estimation

Quality assessment of the raw Illumina sequence data was performed with FastQC v0.11.8 (Andrews et al. 2010). Low quality reads and adapters were removed using Trimmomatic v0.39 (Bolger et al. 2014). The reads were scanned by a 4-based sliding window with average cutting threshold lower than 15 Phred score. Leading and trailing bases were also filtered out with quality score less than 10. Reads with total length shorter than 75 bp and average score below 30 have been omitted.

Adapter trimming and length filtering of basecalled ONT data was done using Porechop v0.2.4 (https://github.com/rrwick/Porechop) with default parameters and the extra option --discard_middle to discard reads with internal adapters.

The genome size was estimated using the k-mer histogram method with Kmergenie v1.7051 (Chikhi and Medvedev 2014) from Illumina data.

### 3. De novo genome assembly

The long ONT reads were used for the construction of a *de novo* assembly, and the Illumina reads were used for the polishing stages. For the initial assembly, we used three different softwares SMARTdenovo (https://github.com/ruanjue/smartdenovo) which produces an assembly from all-vs-all raw read alignments without an error correction stage, Canu v1.8 (Pinto 2014) which relies on the *overlap-layout-consensus* (OLC) method and incorporates an error correction step, and Flye v2.6 (Kolmogorov et al. 2019) algorithm, a repeat graph assembler.

First, we corrected the ONT dataset with Canu, using default parameters except for *corMinCoverage=0*, allowing read correction regardless of the coverage and *corMhapSensitivity=high*, due to the estimated low coverage of our dataset (∼20X). Next, we performed two rounds of assembly, one with SMARTdenovo and one with Canu, with default parameters in both cases. Finally, a third assembly was constructed using Flye with default settings and an approximate genome size of 500 Mb. Based on the quality assessment results (see following section), we decided to proceed with the Flye assembly. We polished the selected assembly with two rounds of Racon v1.4.3 (Vaser et al. 2017), using only preprocessed long reads mapped against the assembly with Minimap2 v2.17 (Li 2018). Further polishing was performed with Medaka v0.9.2 (https://github.com/nanoporetech/medaka) and the final contigs were polished using Pilon v1.23 (Walker et al. 2014) after mapping the Illumina reads against the partially polished assembly with Minimap2 v2.17.

### 4. Quality assessment of draft assemblies

We evaluated our draft assemblies following two methods: (1) the N50 sizes of contigs, using QUAST v5.0.2 (Gurevich et al. 2013), and (2) using BUSCO v3.1.0 (Simão et al. 2015) either standalone or through gVolante (Nishimura et al. 2017) against the Actinopterygii ortholog dataset v9, with default parameters.

The whole pipeline conducted herein is shown in Figure 1.

**Figure 1.**
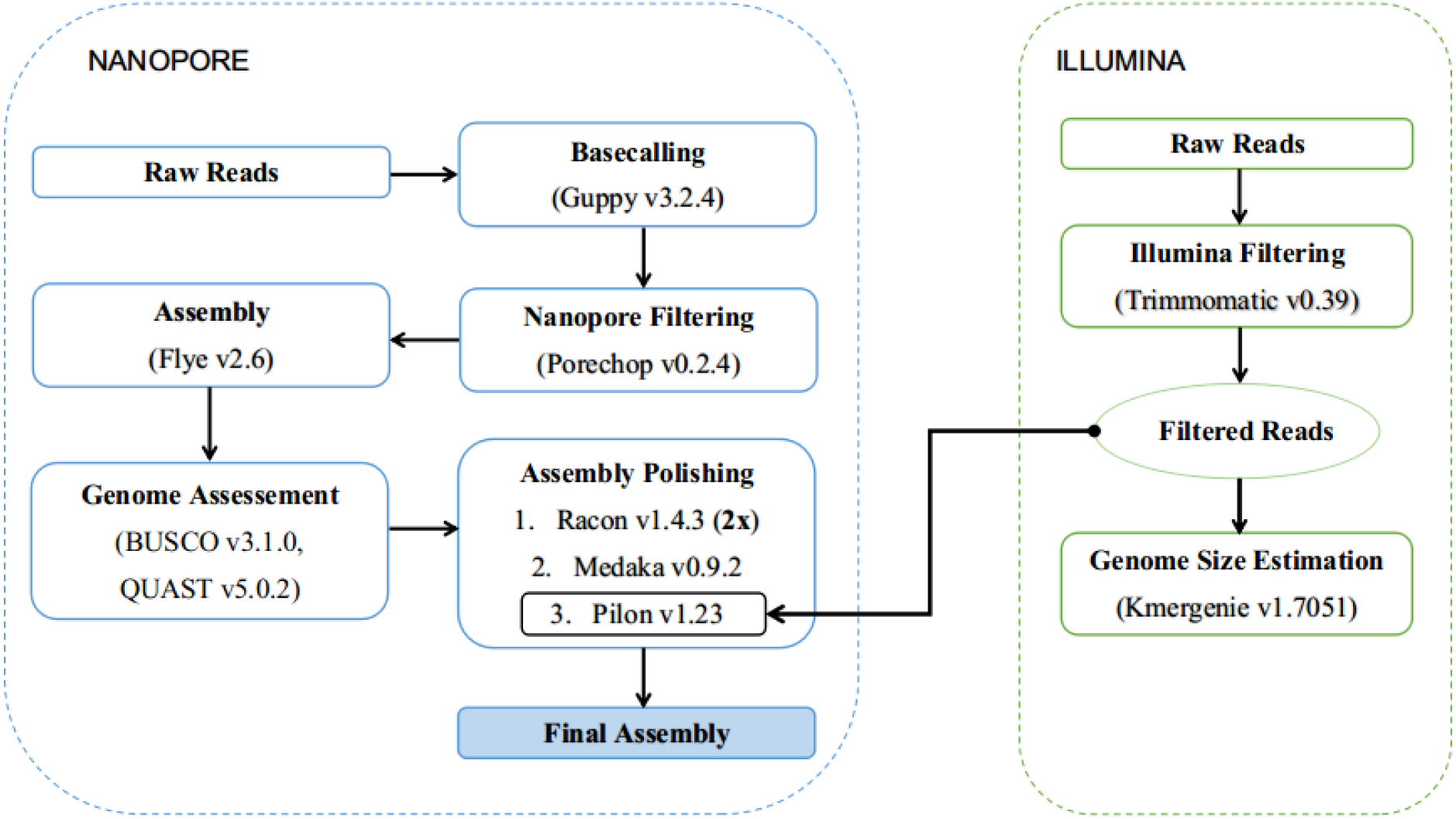
*L. sceleratus* genome assembly pipeline.

## RESULTS & DISCUSSION

Sequencing yielded 17.19 Gb of raw Illumina reads and 9.68 Gb, above Q7, of long ONT reads, with N50 of 48.85 Kb. The estimated genome size was ∼360 Mb and best predicted k = 81. After quality trimming and filtering, we retained 13.34 Gb Illumina data for genome polishing and 9.67 Gb ONT data (Table 1) employed for the genome assembly. The final assembled and polished genome contained 241 contigs with total length of ∼373 Mb, with the largest contig sizing 17 Mb and N50 of 11 Mb (Table 2). The resulted assembly shows that *L. sceleratus* genome size is comparable with that of other puffers, such as *Fugu rubripes* (∼365 Mb; Aparicio et al. 2002), *Takifugu flavidus* (∼377 Mb; Zhou et al. 2019), *Takifugu bimaculatus* (∼393.15 Mb; Zhou et al. 2019), *Takifugu obscurus* (∼373 Mb; Kang et al. 2019) and *Tetraodon nigroviridis* (340 Mb, Jaillon et al. 2004). The contig N50 value (∼11 Mb) of the constructed *L. sceleratus* assembly is considerably greater than that reported for the genomes of *Takifugu bimaculatus* (1,31 Mb; Zhou et al. 2019) and *Takifugu flavidus* (4,4 Mb; Zhou et al. 2019), demonstrating a remarkably contiguous genome assembly. In teleosts, only a few studies, such as in greenfin horse-faced filefish, *Thamnaconus septentrionalis* (22.46 Mb; Bian et al. 2019), red-spotted grouper, *Epimetheus akaara* (5.25 Mb; Ge et al. 2019), two-spotted puffer *Takifugu bimaculatus* (1.31 Mb; Zhou et al., 2019), and yellow-belly puffer *Takifugu flavidus* (4.4 Mb; Zhou et al., 2019) described longer contig N50 than 1 Mb, probably because the main part of their genome assemblies were constructed using ONT or PacBio reads.

**Table 1.** Summary of sequencing results.

| Sequencing technology | Raw Reads | Quality-controlled Reads | Coverage |
| --- | --- | --- | --- |
| Illumina | 57,303,140 | 44,475,382 | 38 x |
| MinION | 552,476 | 484,152 | 20 x |

**Table 2.** Polished genome assembly statistics and completeness.

|  |  |
| --- | --- |
| Total contigs | 241 |
| Total contig sequence | 373,851,781 bp |
| GC (%) | 46.7 |
| Contig N50 | 11,297,640 bp |
| Contig N75 | 6,386,829 bp |
| Longest contig | 17,085,954 bp |
| **BUSCO completeness score** |  |
| Complete | 96.20% |
| Single | 94.40% |
| Duplicated | 2.20% |
| Fragmented | 1.40% |
| Missing | 2.00% |
| Total number of Actinopterygii orthologs | 4,492 (98%) |

According to our knowledge such a highly contiguous reference genome assembly for fish using a single MinION flow cell along with a moderate amount of short Illumina reads has been built only by Bian et al. (2019), who sequenced and assembled the genome of *Thamnaconus septentrionalis*, another member of the order Tetraodontiformes.

Regarding genome completeness, we found 4,513 out of the 4,584 genes, i.e. 98%, of the genes included in the BUSCO Actinopterygian ortholog geneset. Of those, 4,410 (96.20%) were found complete (Table 2), suggesting a high level of completeness and contiguity in the built assembly. Our results are within the same range found in other Tetraodontiodae genomes (e.g. *T. obscurus* [Kang et al. (2019)] and *T. flavidus* [Zhou et al. (2019)]), in spite of not incorporating Hi-C based chromatin data as used in the above referenced studies.

## CONCLUSION

In this study, we present the first highly contiguous and successful genome assembly of *L. sceleratus*. Initially, a primary assembly was constructed with long ONT reads. Polishing with short Illumina reads led to the establishment of a final assembly into contigs with high quality and completeness. These results demonstrate that the Nanopore sequencing method is cost-effective especially for genomes of that size. Since our knowledge about puffer-specific biological aspects is limited, a high-quality *L. sceleratus* genome assembly will enable future comparative studies and investigations on evolutionary and ecological puffer-specific traits. Finally, it will allow further studies on the *L. sceleratus* invasion effectiveness, a unique trait among other Lessepsian migrants.

## ACKNOWLEDGEMENTS

The authors thank Dr. Aspasia Stergioti, Dr. Pantelis Katharios, Chrisa Doxa and Katerina Tasiouli for their assistance in sampling and dissection.

This research was supported through computational resources provided by IMBBC (Institute of Marine Biology, Biotechnology and Aquaculture) of the HCMR (Hellenic Centre for Marine Research). Funding for establishing the IMBBC HPC has been received by the MARBIGEN (EU Regpot) project, LifeWatchGreece RI, and the CMBR (Centre for the study and sustainable exploitation of Marine Biological Resources) RI.

